# Potentiating Na_V_1.1 in Dravet syndrome patient iPSC-derived GABAergic neurons increases neuronal firing frequency and decreases network synchrony

**DOI:** 10.1101/2023.09.28.559990

**Authors:** Matt R. Kelley, Laura B. Chipman, Shoh Asano, Matthew Knott, Samantha T. Howard, Allison P. Berg

## Abstract

Dravet syndrome is a developmental and epileptic encephalopathy characterized by seizures, behavioral abnormalities, developmental deficits, and elevated risk of sudden unexpected death in epilepsy (SUDEP). Most patient cases are caused by *de novo* loss-of-function mutations in the gene *SCN1A*, causing a haploinsufficiency of the alpha subunit of the voltage-gated sodium channel Na_V_1.1. Within the brain, Na_V_1.1 is primarily localized to the axons of inhibitory neurons, and decreased Na_V_1.1 function is hypothesized to reduce GABAergic inhibitory neurotransmission within the brain, driving neuronal network hyperexcitability and subsequent pathology. We have developed a human *in vitro* model of Dravet syndrome using differentiated neurons derived from patient iPSC and enriched for GABA expressing neurons. Neurons were plated on high definition multielectrode arrays (HD-MEAs), permitting recordings from the same cultures over the 7-weeks duration of study at the network, single cell, and subcellular resolution. Using this capability, we characterized the features of axonal morphology and physiology. Neurons developed increased spiking activity and synchronous network bursting. Recordings were processed through a spike sorting pipeline for curation of single unit activity and to assess the effects of pharmacological treatments. At 7-weeks, the application of the GABA_A_R receptor agonist muscimol eliminated network bursting, indicating the presence of GABAergic neurotransmission. To identify the role of Na_V_1.1 on neuronal and network activity, cultures were treated with a dose-response of the Na_V_1.1 potentiator δ-theraphotoxin-Hm1a. This resulted in a strong increase in firing rates of putative GABAergic neurons, an increase in the intraburst firing rate, and eliminated network bursting. These results validate that potentiation of Na_V_1.1 in Dravet patient iPSC-derived neurons results in decreased firing synchrony in neuronal networks through increased GABAergic neuron activity and support the use of human neurons and HD-MEAs as viable high-throughput electrophysiological platform to enable therapeutic discovery.

## Introduction

Dravet syndrome is a rare epileptic encephalopathy with disease onset occurring within the first two years of life. Patients exhibit seizures, episodes of status epilepticus, visual deficits, cognitive delay, age-dependent movement disorders, and elevated risk for premature death (Gataullina & Dulac, 2017). The majority of Dravet syndrome cases result from *de novo* loss-of-function (LOF) mutations in *SCN1A,* the gene coding for the sodium channel protein type 1 subunit alpha, a component of the voltage-gated sodium channel Na_V_1.1. (Dravet, 2011). Genetic mouse models of Na_V_1.1 haploinsufficiency have demonstrated specific reduction of sodium currents and action potential firing in GABAergic inhibitory interneurons as the fundamental cause of pathology resembling Dravet syndrome (Catterall, 2018; Colasante et al., 2020; Yamagata et al., 2020).

Induced pluripotent stem cell (iPSC) technology has enabled the use of human neurons as model systems for studying mechanisms of neurological disorders, including in Dravet syndrome (Takahashi & Yamanaka, 2006). These cellular systems contain identical genetic background to the donor patient, enabling a platform for molecular and physiological translational studies. Advances in cellular programming have resulted in the ability to differentiate many unique neuron types, including neurons with similar features to human forebrain GABAergic interneurons relevant to Dravet syndrome pathophysiology (Liu et al., 2013). For our study, cortical GABAergic neurons were iPSC-derived from a Dravet syndrome pediatric patient with a truncation mutation in *SCN1A.* We electrically characterized the neuronal cultures over 7-weeks using high-density multielectrode arrays (HD-MEAs). Each HD-MEA featured 26,400 electrodes and an electrode pitch of 17.5 μm, enabling recordings at single cell and subcellular resolution (Muller et al., 2015). The neuronal cultures in this study developed spontaneous neuronal firing, neuronal bursting activity, and recurrent, robust network bursting. HD-MEAs also provide a high-throughput and high spatiotemporal resolution platform for obtaining direct electrical readouts of axon potential propagation (Bakkum, Frey, et al., 2013; Emmenegger et al., 2019; Radivojevic et al., 2017). Using HD-MEAs, we recorded from the axons of the Dravet patient iPSC-derived neurons over development, demonstrating the feasibility of this technology for high-throughput recording of axonal physiology features.

Mouse models of Na_V_1.1 haploinsufficiency have supported that prospective therapies which rescue Na_V_1.1 levels should be clinically effective to reduce seizures and ameliorate comorbidities (Han, 2020; Tanenhaus et al., 2022; Valassina et al., 2022; Yamagata et al., 2020). To assess the electrophysiological effects of activating Na_V_1.1 in the Dravet syndrome patient iPSC-derived GABAergic neurons, we treated the neuronal cultures with Hm1a, a spider venom peptide that selectively potentiates Na_V_1.1 (Osteen et al., 2016).

Intracerebroventricular infusion of Hm1a in a mouse Dravet syndrome model of Na_V_1.1 haploinsufficiency has been previously demonstrated to rescue action potential firing in GABAergic interneurons without affecting excitatory neuronal firing, resulting in reduction of seizure count and mortality (Mattis et al., 2022; Richards et al., 2018). We found that Hm1a increased the firing rate and intraburst frequency of the human GABAergic neurons, along with decreasing neuronal network synchrony. These results support HD-MEA technology combined with patient iPSC-derived neurons as a high throughput platform for drug discovery research and targeting rescue of Na_V_1.1 function in Dravet syndrome.

## Results

### Development of human iPSC-derived neuron *in vitro* model of Dravet syndrome

An induced pluripotent stem cell (iPSC) derived from a Dravet syndrome patient was differentiated into a population of enriched cortical GABAergic neurons (Fig1 1a). The patient possessed a frameshift mutation in *SCN1A* resulting in a truncation. The identity of neurons expressing GABA was confirmed by immunostaining at DIV 7 for the neuronal marker MAP2 and GABA. At this timepoint, 69% of cells were positive for the neuronal marker MAP2, and 64% of MAP2 cells were positive for GABA (Figure 1b). To understand the relative expression of *SCN1A* across time in culture, a RT-qPCR experiment was performed from RNA extracted from days *in vitro* (DIV) −10, −14, −21, and −28. The fold change of *SCN1A* expression for each timepoint to DIV 10 was determined (mean fold change ± SD. DIV-14: 1.3 ± 0.5, DIV-21: 1.9 ± 0.1, *P* < 0.01, DIV-28: 2.7 ± 0.2, *P* < 0.0001, *n*=8 biological replicates per timepoint). This demonstrated an upregulation of *SCN1A* expression across 4 weeks *in vitro* in the neuronal cultures.

**Figure 1.**
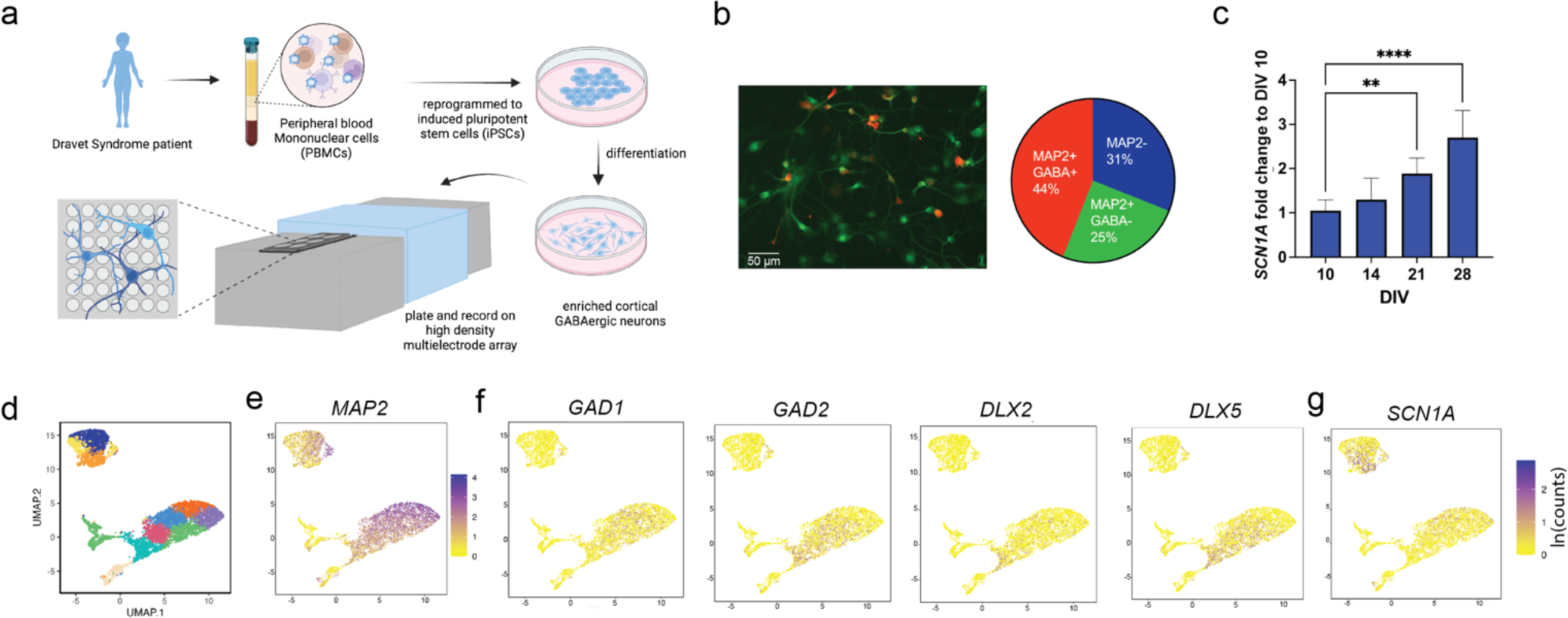
Dravet syndrome patient-derived iPSC Cortical GABAergic neurons upregulate *SCN1A* and are positive for neuronal markers. a) Cortical GABAergic neurons were produced from Dravet syndrome patient-derived iPSCs and activity assessed using high density multielectrode arrays. b) Neurons were plated and immunostained for the neuronal marker MAP2 and GABA, 44% of cells were positive for both markers. c) Quantitative PCR on RNA extracted from neuronal cultures demonstrated an upregulation of *SCN1A.* d) Single cell RNA sequencing of the neurons performed at DIV 21 mapped to 11 clusters. e-f) Cells demonstrated strong expression of neuronal and GABAergic marker genes. e) Cells broadly expressed the neuronal marker MAP2. f) Cells expressed multiple markers of GABAergic neurons along with g) *SCN1A.* ** *P* < 0.01, **** *P* < 0.0001. One-way ANOVA.

### Characterization of Dravet syndrome patient iPSC-derived neurons by single cell RNA sequencing

To further characterize the neurons, a single cell RNA sequencing experiment was performed from neurons dissociated at DIV 21. The transcriptomes of 6208 cells were sequenced and after quality control and preprocessing, single cells were grouped off gene expression profiling and visualized using uniform manifold approximation and projection (UMAP) plots (Fig 1d). To characterize these cells, we looked at which cells were expressing neuronal and GABAergic markers. The neuronal marker, *MAP2*, was detected in 90.2% of cells while 42.3% of sequenced cells expressed at least one GABAergic marker (Fig 1e-f). As expected, most cells, (99.7%), expressing GABAergic marker transcripts were also positive for *MAP2*. Given we were looking at Dravet patient-derived neurons, we assessed the expression patterns of the disease-causing gene, *SCN1A* which was expressed in 38.1% of cells (Fig 1g). Of the 2366 cells expressing *SCN1A*, 46.3% expressed at least one of the GABAergic markers: *GAD1*, *GAD2*, *DLX2*, or *DLX5*. While the cultures were enriched for GABAergic neurons as expected, a minority of cells expressed other neuronal cell type markers, such as 12% of cells that expressed the glutamatergic marker *SLC17A6* (data not shown). Of the 746 *SLC17A6* expressing cells, 399 were positive of *SCN1A.* These scRNA-seq results demonstrate that the iPSC patient-derived neurons were enriched for GABAergic neuronal markers and a large population of these neurons expressed *SCN1A*.

### Dravet syndrome patient iPSC-derived neuronal cultures on HD-MEAs exhibited neuronal firing and network synchrony

We used HD-MEAs to record electrophysiological activity from the Dravet syndrome patient iPSC-derived cortical GABAergic neurons and characterized the neuronal cultures through development. The neurons were plated with apparently normal human iPSC-derived astrocytes to support viability of the cultures over the 7-week time course of the experiment. The arrays were fixed at end of experiment at DIV 42, labeled with GAD-67 and VGLUT2 antibodies and imaged by confocal microscope. As the cultures developed, these microscopy images confirmed that dense neuronal networks formed on the array surface (Fig 2a). To map electrical activity of each culture, an activity scan of electrodes was performed across each HD-MEA surface at weekly timepoints. All *n*=12 HD-MEAs plated for this study were electrically active and exhibited spontaneous neuronal firing well above the spike detection threshold of 5.5 standard deviations of the average voltage of recording traces (Figure 2b). Through analysis of the activity scan data, electrical images were constructed to depict the activity of the cultures in firing rate and spike amplitude over the 7-week time course (Fig 2c). Observed activity increased over time in percent of channels active across the array, mean channel firing rate, mean channel interspike interval and mean channel spike amplitude. (Fig 2d-g. *n*=12 HD-MEAs, DIV 14 to DIV 42: Active channels: 13.0 ± 1.6% to 42.5 ± 3.7%, firing rate: 0.57 ± 0.02 Hz to 1.31 ± 0.05 Hz, interspike interval: 85.7 ± 1.12 ms to 101.7 ± 0.95 ms, *P* < 0.0001, spike amplitude: 36.4 ± 0.3 µV to 40.0 ± µV, *P* < 0.001). Using the activity scan data, a network recording was performed in the neuronal-unit configuration (see methods) with 1020 channels recorded in parallel. Starting at DIV 28, the neuronal cultures showed network bursting. Bursting events were detected by the ISI_N_ algorithm, to determine burst start and stop timepoints (Bakkum, Radivojevic, et al., 2013). At DIV 42, networks showed robust and recurring bursting events (Figure 2h). The bursts featured a rise in instantaneous average firing frequency of the channels followed by a depression in firing rate (Figure 2i). There was an increase of burst duration, intraburst peak firing rate, interburst interval, the number of spikes per burst, and a decrease in burst interspike interval, from DIV 28 to 42 (Figure 2j-m, Table 1).

**Figure 2.**
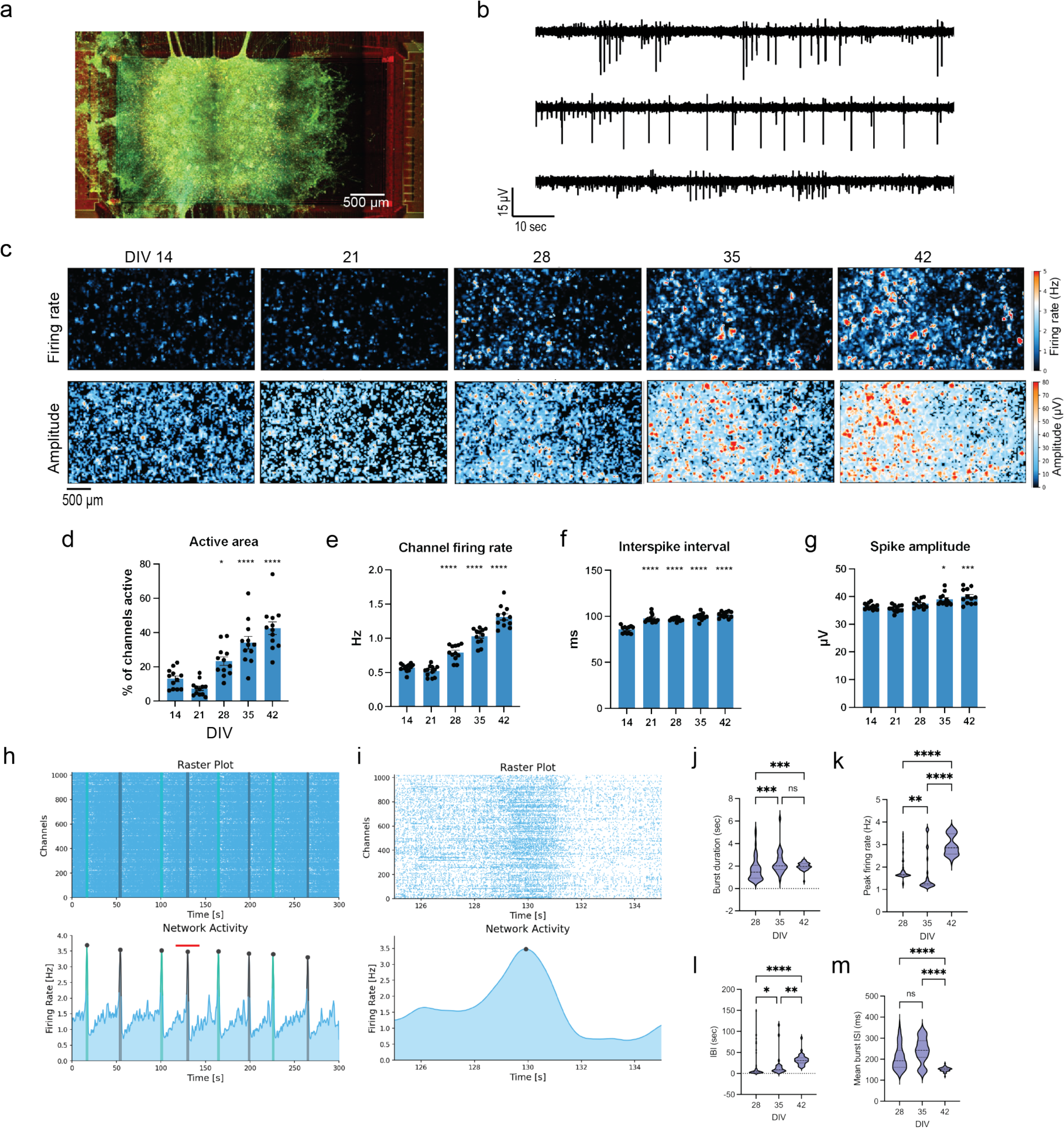
Dravet syndrome patient iPSC-derived neurons fire spontaneously and display network bursting when plated on HD-MEAs. a) Following plating of neurons and astrocytes on HD-MEAs, recordings were performed weekly till DIV 42. Representative image of a HD-MEA well at DIV 42, with GAD-67 (green) VGLUT2 (yellow). b) HD-MEAs used permitted recording channels from 1020 electrodes in parallel. Three example traces from individual channels. c) Each HD-MEA was scanned weekly for firing activity using the Activity Scan assay (Maxwell Biosystems). Images of a HD-MEA well across time in culture showing firing rate (top row) and spike amplitude (bottom row) of scanned channels. d-g) Features of the neuronal cultures derived from activity scans of *n*=12 HD-MEAs. Each point represents the average from 1 MEA. Statistics are means compared to DIV 14. From DIV 14 to 42, there was an increase in HD-MEA active area, channel firing rate, interspike interval, and spike amplitude. h) Cultures plated on the HD-MEAs displayed synchronous network bursting, detected by ISI_N_ algorithm. Raster plot of a 5-minute recording at DIV 42 plotted above the average instantaneous spike frequency of channels. Each detected burst is highlighted by the grey and green shading, with black dot identifying peak. i) Magnified raster plot and instantaneous firing frequency of burst identified by red line in e. j - m) Features of network bursts detected from bursting events from DIV 28, 35, and 42. There was an increase in burst duration, intraburst peak firing rate of the average instantaneous spike frequency, IBI, and decrease in the mean burst ISI. ns: no significant difference, * *P* <0.05, ** *P* < 0.01, *** *P* < 0.001, **** *P* < 0.0001. Kruskal-Wallis test, followed by the Dunn-Sidák multiple-comparison test.

**Table 1.** Burst features of Dravet syndrome patient-derived iPSC Cortical GABAergic neurons.

|  | Days *in vitro* |  |  |  |  |  |
| --- | --- | --- | --- | --- | --- | --- |
|  | 28 | 35 | 42 | DIV 28 to 35 | DIV 28 to 42 | DIV 35 to 42 |
| **Burst event count** | 81 | 31 | 49 |  |  |  |
| **Burst features (mean ± SEM)** |  |  |  |  |  |  |
| Burst duration (sec) | 1.72 ± 0.12 | 2.29 ± 0.18 | 2.00 ± 0.05 | *** | *** | ns |
| Intraburst peak firing rate (Hz) | 1.85 ± 0.05 | 1.62 ± 0.14 | 2.97 ± 0.07 | ** | **** | **** |
| Interburst interval (sec) | 14.0 ± 2.4 | 17.9 ± 5.1 | 32.7 ± 2.1 | * | **** | ** |
| Mean burst interspike interval (ms) | 208.2 ± 6.5 | 241.0 ± 10.6 | 151.8 ± 1.5 | ns | **** | **** |
| Number of spikes in burst | 3161 ± 237 | 2640 ± 399 | 5396 ± 179 | ns | **** | **** |

Kruskal-Wallis test, followed by the Dunn-Sidák multiple-comparison test. \* $P < 0.05$ , \*\* $P < 0.01$ , \*\*\* $P < 0.001$ , \*\*\*\* $P < 0.0001$

### Characterization of axonal morphology and physiology in Dravet syndrome patient iPSC-derived neurons using HD-MEAs

HD-MEA technology enables recording neuronal activity at subcellular resolution and a non-invasive method to record axon potential propagation (Bakkum, Frey, et al., 2013; Emmenegger et al., 2019; Radivojevic et al., 2017). In this study, we used this capability of HD-MEAs to characterize the development of Dravet syndrome patient iPSC-derived neuronal axons over development *in vitro* and measured features of axonal physiology. *N*=6 HD-MEAs were recorded weekly through DIV 42. Following a scan for neuronal activity, thirty putative neurons were identified on each HD-MEA using a spike sorting algorithm included with the AxonTracking assay (Maxwell Biosystems AG). 1532 neurons were identified from this experiment, and features of axons characterized (Table 2). Tracking of axons across the surface of the HD-MEAs showed widespread axonal arbors (Figure 3a). Neurons demonstrated a variety of axon shapes and lengths that were electrically imaged on the MEA surface through a heat map of the maximum detected voltage trace of each axon (Figure 3b). Recordings from electrodes along the axon enabled derivation of the axon potential conduction velocity for each neuron (Figure 3c). There as an increase over DIV 35 to DIV 42 in neuron firing rate, amplitude at the axon initiation site, and total axon lengths (Figure 3d - 3f). The conduction velocity, longest branch length, and longest latency of spike from initiation site to terminal synapse plateaued prior to DIV 42 (Figure 3g – 3i). These results confirmed the maturations of the Dravet patient iPSC-derived neurons and the feasibility of HD-MEA technology for high-throughput recording of axonal physiology features.

**Table 2.** Features of Dravet syndrome patient-derived iPSC Cortical GABAergic neurons.

|  | Days *in vitro* |  |  |  |  |  |  |  |  |  |
| --- | --- | --- | --- | --- | --- | --- | --- | --- | --- | --- |
|  | 21 | 28 | 35 | 42 |  |  |  |  |  |  |
| Neuron count | 199 | 367 | 475 | 491 |  |  |  |  |  |  |
| Axon features (mean ± SEM) |  |  |  |  | DIV 21 to 28 | DIV 21 to 35 | DIV 21 to 42 | DIV 28 to 35 | DIV 28 to 42 | DIV 35 to 42 |
| Firing rate (Hz) | 1.45 ± 0.05 | 2.31 ± 0.07 | 3.00 ± 0.07 | 3.33 ± 0.08 | **** | **** | **** | **** | **** | ns |
| Amplitude at initiation site (µV) | 16.8 ± 0.3 | 19.2 ± 0.4 | 22.5 ± 0.5 | 22.9 ± 0.5 | **** | **** | **** | **** | **** | ns |
| Total axon length (mm) | 0.91 ± 0.08 | 1.77 ± 0.12 | 2.31 ± 0.11 | 3.29 ± 0.13 | **** | **** | **** | **** | **** | **** |
| Conduction velocity (m/s) | 0.484 ± 0.006 | 0.506 ± 0.004 | 0.513 ± 0.004 | 0.521 ± 0.003 | *** | **** | **** | ns | ns | ns |
| Longest branch length (mm) | 0.40 ± 0.02 | 0.50 ± 0.02 | 0.53 ± 0.02 | 0.56 ± 0.02 | * | **** | **** | ns | **** | ns |
| Longest latency (ms) | 2.1 ± 0.1 | 2.8 ± 0.1 | 3.3 ± 0.1 | 3.5 ± 0.1 | *** | **** | **** | *** | **** | ns |

Kruskal-Wallis test, followed by the Dunn-Sidák multiple-comparison test. \* *P* < 0.05, \*\* *P* < 0.01, \*\*\* *P* < 0.001, \*\*\*\* *P* < 0.0001

**Figure 3.**
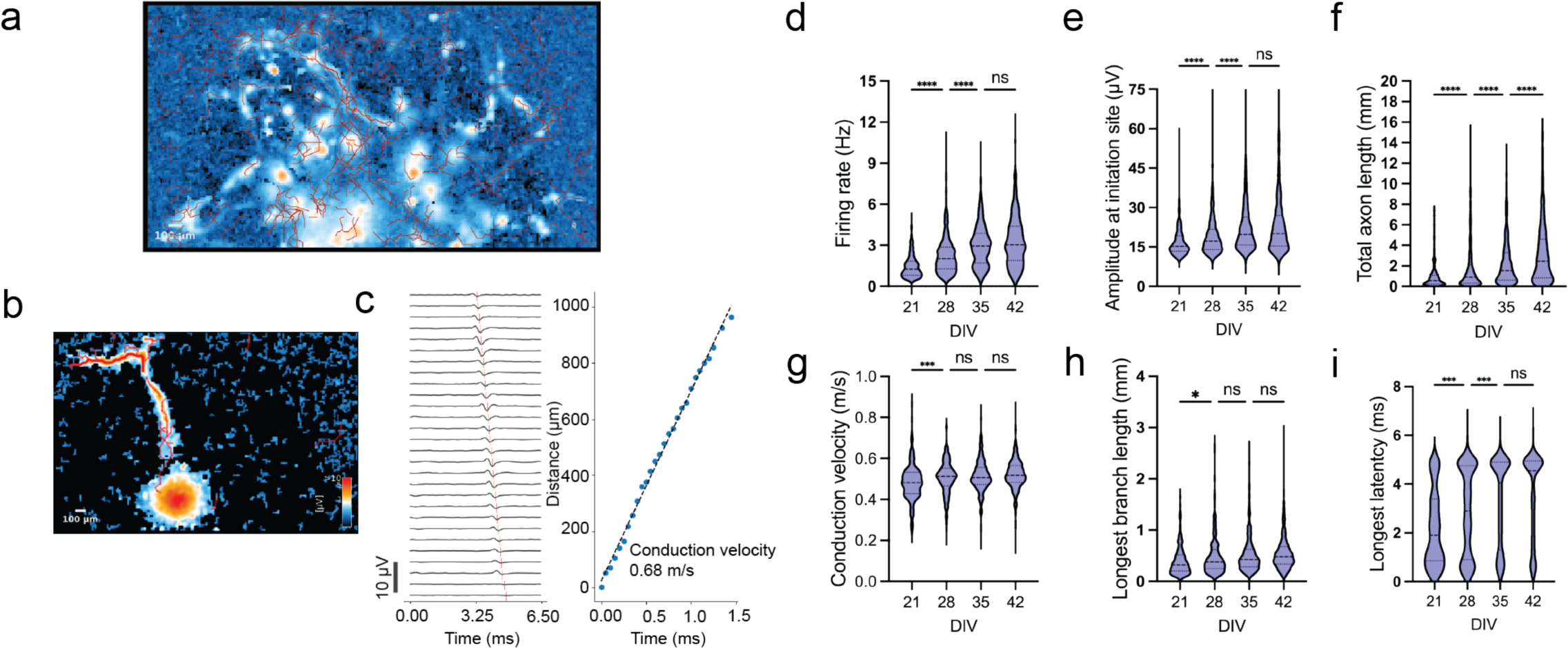
High-density MEA recordings enabled detection and characterization of Dravet patient iPSC-derived neuronal axons. 1844 neurons were recorded for this experiment. a) Map of a HD-MEA with reconstructed axons (red LINES) detected using the AxonTracking Assay (Maxwell Biosytems). b) An example identified neuron with detected axon branches traced in red. c) Raw traces of detected axon extracellular action potential waveform (EAP) from neuron branch indicated in b by bold red permitted calculation of conduction velocity over 1 mm distance. d-i) Features of detected neurons and axons from week 3 to 6. ns: no significant difference, * *P* <0.05, *** *P* < 0.001, **** *P* < 0.0001. Kruskal-Wallis test, followed by the Dunn-Sidák multiple-comparison test.

### Dravet syndrome patient iPSC-derived neurons displayed a diversity of neuronal firing phenotypes

Following an activity scan for neuronal activity on each plated HD-MEA, select electrodes were routed for a neuronal network recording from putative neuronal units. Spike-sorting was performed on each preprocessed 5-minute recording from *n*=6 HD-MEAs followed from DIV 28, 35 and 42. This facilitated characterization of individual neurons and neuronal network features of the Dravet syndrome iPSC-derived neurons (Figure 4a, features in Table 3). An average of 366 ± 25 neurons was sorted per HD-MEA, with *n*=6 HD-MEAs per timepoint. The average maximum voltage extracellular action potential (EAP) waveform was used for waveform feature extraction and identifying the location of neurons on the HD-MEA surface (Figure 4b). Waveform features were extracted from each neuron (Figure 4d, Table 3). Neurons displayed various firing modes, including regular intervals of spiking or discrete bursting episodes (Figure 4e). Analysis of the extracted waveform features showed a change through development. From DIV 28 to 42, there was a decrease in mean spike amplitude, half-width, peak latency, asymmetry ratio, decay, recovery slope, and an increase in trough-to-peak ratio, repolarization slope (Figure 4f). Unit activity features identified the firing and bursting properties of the neurons. In analysis of the complete cohort of spike sorted neurons, there was an increase in firing rate over development (Figure 4f). Neurons were assessed for bursting activity by the logISI method (Pasquale et al., 2010) (Figure 4g). There was a limited change in the percent of neurons at each DIV timepoint characterized as bursting (Figure 4h. DIV-28: 46.1%, DIV-35: 50.7 %, DIV-42: 49.1%) This showed that across development from 28 to 42 DIV, the percent of neurons with bursting activity remained consistent. From DIV 28 to 42, there was no significant change in the mean burst frequency of occurrence or burst duration (Figure 4i). Identified bursting neurons demonstrated mean intraburst frequencies near 30 Hz with some neurons firing at intraburst frequency of over 40 Hz (Figure 4i). Intraburst firing rate increased over development from DIV 28 to 42. For each neuronal culture, the spike timepoints were assessed by the ISI_N_ algorithm (Bakkum, Radivojevic, et al., 2013) for neuronal network bursting activity. Repeat and robust network bursting events were detected on all cultures (Figure 4j). The frequency of network bursts and burst duration increased over development from DIV 28 to 42. However, there was no significant difference in number of neuronal spikes per burst over the similar period. These results showed that HD-MEA recordings provide a platform for high throughput characterization of neuron and neuronal network features across multiple weeks of development *in vitro*.

**Figure 4.**
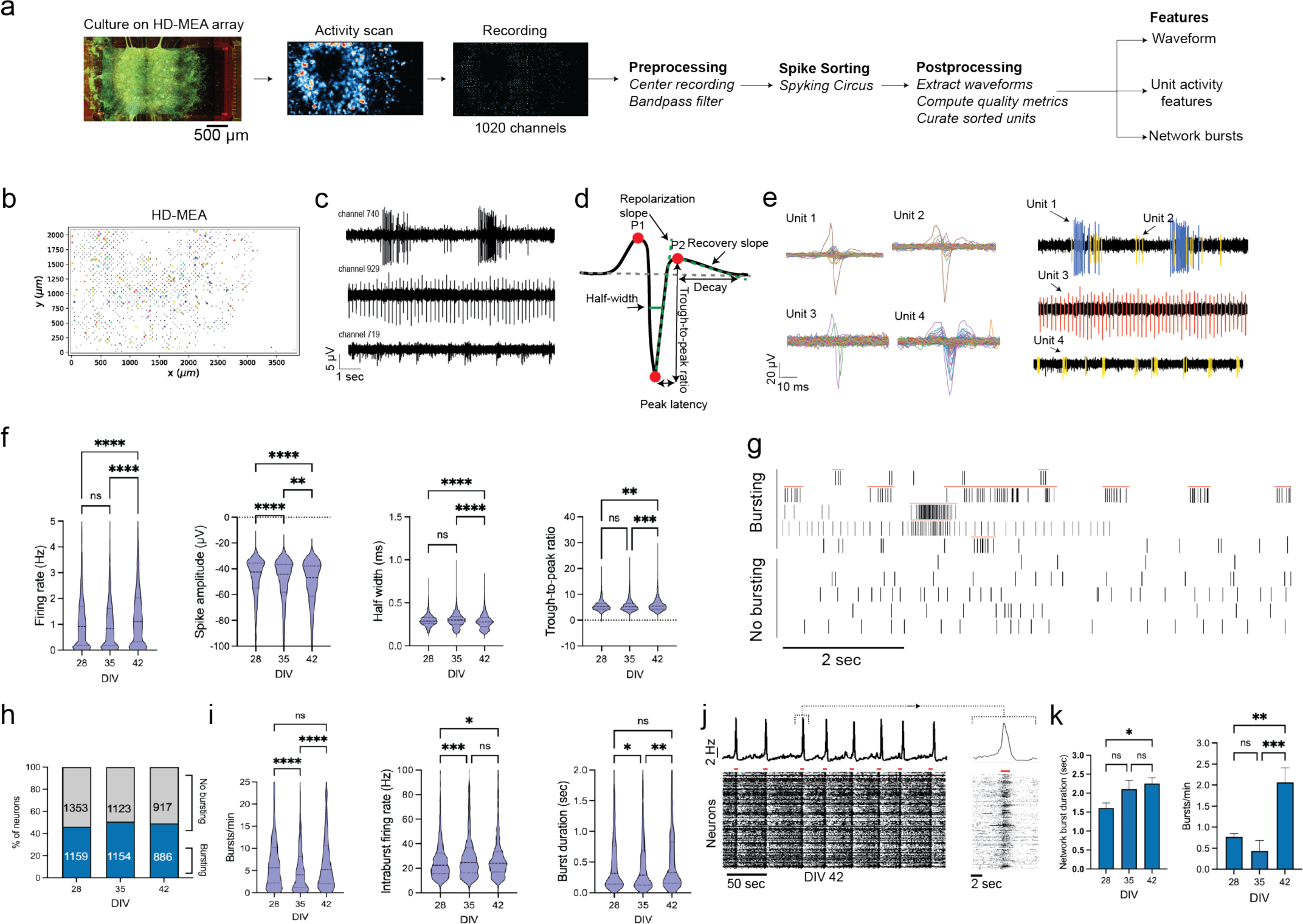
Spike sorting was performed on HD-MEA recorded Dravet syndrome patient iPSC derived neuronal cultures to identify and characterize neuronal features and contribution to network activity. a) Recordings were preprocessed and spike sorting performed using the SpykingCircus algorithm. Features were extracted from the extracellular action potential waveform (EAP), neuronal unit and network activity. b) Map of a HD-MEA probe and recorded unit locations. Grey squares indicated recorded 1020 electrodes and colored dots the location of detected max waveforms of spike sorted units. c) Recordings from indicated HD-MEA channels. d) Depiction of features extracted from average maximum voltage EAP of each spike sorted neurons. e) Average waveforms of 4 units detected from channels shown in c. Neuron EAPs overlayed on recordings. f) Selected features extracted from neuronal waveforms. There was an observed increase in neuron firing rate, spike amplitude, half width and trough-to peak ratio over development. g) Raster plot of 10 units over a ten second window, each marker represents neuronal spike. Red lines indicate detected bursting activity by ISI_N_ algorithm. h) Percentage of neurons with bursting activity at DIV 28, 35, and 42. i) Neurons displayed an increase of burst frequency, intraburst firing rate, and burst duration over development. j) The instantaneous spike frequency plotted over a raster plot of neuron spike events from an HD-MEA at DIV 42. Cultures displayed synchrony and robust network bursting k) Network burst duration and frequency increased over development. ns: no significant difference, * *P* <0.05, ** *P* < 0.01, *** *P* < 0.001, **** P < 0.0001. Kruskal-Wallis test, followed by the Dunn-Sidák multiple-comparison test.

**Table 3.** Waveform, unit activity and network activity of Dravet syndrome patient-derived iPSC cortical GABAergic neurons.

|  | Days *in vitro* |  |  |  |  |  |
| --- | --- | --- | --- | --- | --- | --- |
|  | 28 | 35 | 42 | DIV 28 to 35 | DIV 28 to 42 | DIV 35 to 42 |
| **Neuron count** | 2512 | 2277 | 1803 |  |  |  |
| **Waveform features (mean ± SEM)** |  |  |  |  |  |  |
| Amplitude (μV) | -47.6 ± 0.3 | -50.7 ± 0.4 | -52.6 ± 0.5 | **** | **** | *** |
| Half width (ms) | 0.29 ± 0.00 | 0.3 ± 0.00 | 0.28 ± 0.00 | ns | **** | **** |
| Peak latency (ms) | 0.76 ± 0.01 | 0.75 ± 0.01 | 0.74 ± 0.01 | ns | **** | * |
| Asymmetry ratio | -0.26 ± 0.01 | -0.29 ± 0.01 | -0.28 ± 0.01 | **** | **** | * |
| Trough to peak ratio | 5.61 ± 0.04 | 5.69 ± 0.05 | 5.98 ± 0.06 | ns | ** | *** |
| Decay (ms) | 4.65 ± 0.01 | 4.64 ± 0.01 | 4.61 ± 0.01 | ns | ** | *** |
| Repolarization slope | 19.4 ± 0.2 | 21.3 ± 0.3 | 23.5 ± 0.4 | ** | **** | *** |
| Recovery slope | -1.2 ± 0.1 | -1.3 ± 0.0 | -1.3 ± 0.0 | **** | **** | ns |
| **Unit activity features (mean ± SEM)** |  |  |  |  |  |  |
| Spike frequency (Hz) | 1.2 ± 0.0 | 1.1 ± 0.0 | 1.5 ± 0.03 | ns | **** | **** |
| Interspike interval (ms) | 3.66 ± 0.09 | 3.63 ± 0.10 | 3.29 ± 0.13 | ns | **** | **** |
| Intraburst frequency (Hz) | 27.7 ± 0.7 | 33.7 ± 1.4 | 32.5 ± 1.5 | *** | * | ns |
| Burst frequency (bursts/min) | 7.6 ± 0.2 | 6.2 ± 0.2 | 8.3 ± 0.3 | **** | ns | **** |
| Burst duration (sec) | 0.67 ± 0.03 | 0.53 ± 0.02 | 0.57 ± 0.02 | * | ns | ** |
| Number of spikes in burst | 9.99 ± 0.41 | 8.62 ± 0.27 | 9.50 ± 0.30 | ns | * | *** |
| Intraburst interspike interval (ms) | 75.3 ± 2.2 | 69.8 ± 2.0 | 70.2 ± 2.2 | ns | ns | ns |
| **Network activity features (mean ± SEM)** |  |  |  |  |  |  |
| Burst frequency (bursts/min) | 0.8 ± 0.1 | 0.4 ± 0.3 | 2.1 ± 0.4 | ns | ** | *** |
| Burst duration (sec) | 1.31 ± 0.13 | 2.11 ± 0.23 | 2.25 ± 0.15 | ns | * | ns |
| Spikes per burst | 2297 ± 186.1 | 4323 ± 666.0 | 3003 ± 174.2 | ** | ns | ns |

Kruskal-Wallis test, followed by the Dunn-Sidak multiple-comparison test. \* $P < 0.05$ , \*\* $P < 0.01$ , \*\*\* $P < 0.001$ , \*\*\*\* $P < 0.0001$

### Potentiation of Na_V_1.1 by the peptide Hm1a decreased network synchrony while increasing neuronal firing rate and neuronal intraburst spike frequency

Hm1a is a selective peptide potentiator of Na_V_1.1 derived from spider venom, shown in a mouse Dravet syndrome model of NaV1.1 haploinsufficiency to rescue action potential firing in GABAergic interneurons without affecting excitatory neuronal firing (Osteen et al., 2016; Richards et al., 2018). We treated the neuronal cultures on HD-MEAs at DIV-42 with Hm1a to assess the physiological effect of Na_V_1.1 potentiation on neuron and neural network activity. Along with Hm1a, the GABA_A_ receptor agonist muscimol treatment was performed to assess for functional GABA_A_R mediated signaling. Treatments were added directly into the cell culture media to dilute to the indicated final concentration (Figure 5a). Following an activity scan for neuronal activity on each plated HD-MEA, select electrodes were routed for a neuronal network recording from putative neuronal units. After a baseline recording period, 4 treatments were performed: PBS (vehicle), muscimol (10 μm), Hm1a (100 nM and 500 nM), *n*=3 HD-MEAs per treatment. These recordings were spike sorted as above to identify individual neurons and neuronal activity features used to assess treatment effects on neuron firing and neuronal network synchrony. Application of the GABA_A_ receptor agonist, muscimol, to the neuronal cultures eliminated the occurrence of recurring network bursts that were observed on the cultures at DIV 42 and during the baseline period prior to drug treatment (Figure 5b). Application of Hm1a at 100 nM and 500 nM resulted in a rise in average instantaneous mean firing rate followed by a return to baseline activity and elimination of further network bursts (Figure 5b). To assess the effects of the treatments on the individual neuron firing rate, a 30 second baseline period was compared with 30 second period initiated at the start of the treatment added to the culture media. Each 30 second period was split into 5 second bins of averaged firing rate. The maximum firing rate was compared for each neuron between baseline and treatment to determine a Δ frequency in hertz. The vehicle PBS resulted in a mean Δ frequency of −0.02 ± 0.03 Hz. (Figure 5e). Muscimol resulted in mean Δ frequency of −0.00 ± 0.11 Hz (Figure 5f). Hm1a (100 nM) resulted in mean Δ frequency of 3.51 ± 0.31 Hz and Hm1a (500 nM) resulted in a Δ frequency of 4.35 ± 0.36 Hz (Figure 5g). Hm1a treatment resulted in an increase of Δ frequency when compared to PBS treated neurons, but there was no significant difference in Δ frequency between 100 and 500 nM concentrations. Neurons were assessed for unit bursting activity by the LogISI algorithm (Pasquale et al., 2010). For the 30 second baseline and treatment periods, Hm1a (100 nM) resulted in an increase in percent of neurons classified as bursting (baseline 23%, *n*= 78 neurons, versus Hm1a: 53%, *n*= 178 neurons). There was no significant difference in burst duration due to Hm1a (baseline: 0.51 ± 0.13 sec, Hm1a: 1.25 ± 0.28, *P* = 0.917). Hm1a resulted in an increase of the average number of spikes per neuron per burst (baseline: 7.92 ± 1.78 spikes, Hm1a: 17.12 ± 3.35 spikes, *P* = 0.03). Hm1a increased the mean neuron intraburst firing frequency (baseline: 33.4 ± 3.2 Hz, Hm1a: 43.6 ± 2.6 Hz, *P* < 0.01).

**Figure 5.**
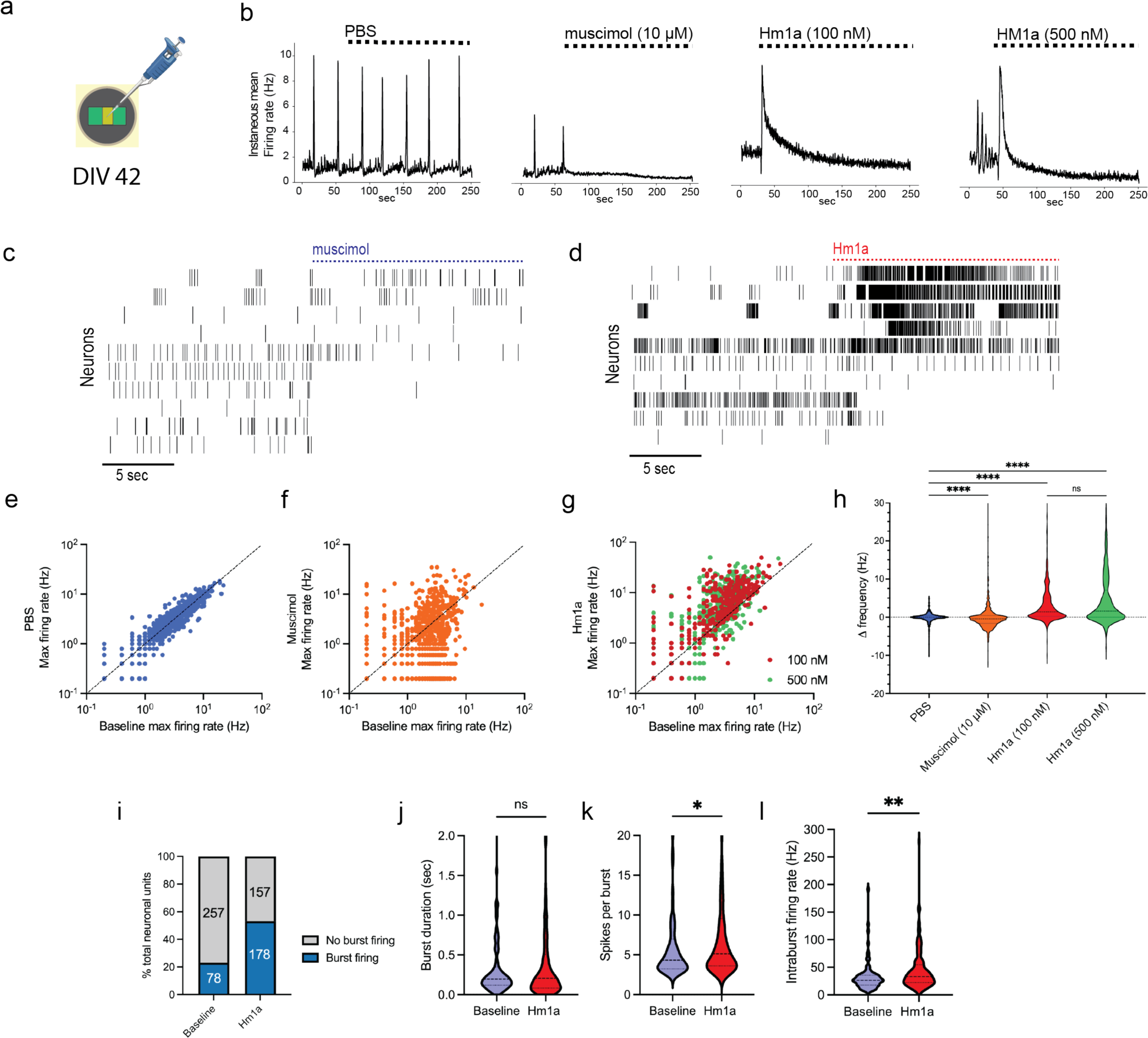
The peptide Hm1a decreased network synchrony and increased the firing of Dravet patient iPSC-derived enriched GABAergic neurons. a) At DIV 42, cultures were treated with drug or vehicle by pipette into cell culture media to achieve indicated concentrations. b) Plots of instantaneous spike frequency of cultures before and after treatments. Both muscimol and Hm1a halted the occurrence of recurring network bursts. c) Raster plot of spike events of 10 neurons on application of muscimol, each marker represents neuronal spike. d) Raster plot of spike events of 10 neurons on Hm1a (100 nM), each marker represents neuronal spike. Hm1a resulted in a dramatic increase of firing of multiple units and reduction of firing of others. e-g) Plots depicting neurons firing rate pre and post indicated treatment. The firing rate was calculated in 5 seconds bins over 30 seconds baseline and 30 seconds post treatment. h) The Δ frequency (Hz) was calculated as difference between treatment firing rate and baseline firing rate. Hm1a results in an increase in neuronal firing. i) Hm1a (100 nm) treatment resulted in an increase of percentage of neurons with detected bursting activity. Numbers in bar graph are total count of neurons. j) Mean burst duration of neurons 30 seconds before and 30 seconds after Hm1a (100 nM) treatment. k) Mean number of spikes per burst before and after Hm1a (100 nM) treatment. l) Neuron intraburst firing rate before and after Hm1a (100 nM) treatment. ns: no significant difference, * *P* <0.05, ** *P* < 0.01, **** *P* < 0.0001. Kruskal-Wallis test, followed by the Dunn-Sidák multiple-comparison test.

## Discussion

In this study, we assessed the effects of Na_V_1.1 potentiation in Dravet syndrome patient iPSC-derived GABAergic neurons, using the high-throughput electrophysiology recording capability of HD-MEA technology. Neurons were maintained and recorded for 7-weeks on the arrays, demonstrating the robustness of this approach to track the development of human derived neuron cultures *in vitro*. Neurons were spontaneously active and increased in activity over the experiment, forming axonal networks, firing in bursts, and contributing to neuronal network synchronous bursting. The HD-MEA platform enabled high-fidelity recording from axons over development, and we obtained measurements of morphology and axon potential electrophysiological features of 1532 neurons in this study. Treatment of the neurons with the Na_V_1.1 potentiator Hm1a increased average neuronal firing rate, neuronal intraburst spike frequency, and eliminated the occurrence of synchronous network bursting.

Human iPSC-derived neurons have emerged as valuable models in enabling disease biology research and drug discovery in neurological disorders, including epilepsy and Dravet syndrome (Simkin et al., 2022). Studies using Dravet syndrome patient iPSC-derived GABAergic neurons have impaired sodium current activation, decreased actional potential firing, hyperactive neuronal networks, and dysregulated pathways for chromatin remodeling (Liu et al., 2016; Schuster et al., 2019; Sun et al., 2016; Xie et al., 2020). A limitation of our study is the lack of isogenic control iPSC-derived neuron line or genetic correction. However, the patient line we used here displayed high spontaneous activity and network bursting, permitting the pharmacology experiments to assess single neuron and neuronal network responses to Na_V_1.1 potentiation. Interneurons compose around 30% of neurons in the human cortex, compared to around 12% of neurons in the mouse cortex (Loomba et al., 2022). Compared with mouse models, human iPSC-derived neuron models of Dravet syndrome can contain a percentage of interneurons closer to that of the human brain, enabling translational models of neuronal networks and determination of contribution of deficits in inhibitory versus excitatory neurons. In our study, the single cell RNA transcriptome confirmed that the 42.3% of cells sequenced in this study expressed at least one GABAergic neuron marker. Other Dravet syndrome iPSC-derived human neuron models with percentages of interneurons in the neuronal network similar to that of the human brain support that Dravet syndrome pathophysiology derives from defective inhibitory neurons (Higurashi et al., 2013; Liu et al., 2016; Sun et al., 2016).

Treatment of the Dravet patient iPSC-derived neurons with the Na_V_1.1 potentiator Hm1a increased firing rate, the percentage of neurons with bursting activity, and the neuronal intraburst firing frequency. Hm1a is a highly selective potentiator of Na_V_1.1, delaying channel inactivation (Osteen et al., 2017). In mice carrying the human Dravet syndrome *Scn1a*^+/R1407X^ variant, Hm1a treatment had minimal effect on firing properties of excitatory CA1 pyramidal neurons while rescuing action potential firing collapse of inhibitory CA1 GABAergic interneurons (Richards et al., 2018). A similar finding was found in the *Scn1a*^+/−^ mouse dentate gyrus, where Hm1a rescued the fast-spiking discharges of parvalbumin positive interneurons and had no effect on evoked granule cell activation. In our experiment, the majority of Dravet syndrome patient iPSC-derived neurons showed an increase in firing rate due to Hm1a treatment and there was an observed increase in the mean intraburst firing frequency of the neurons from 33.4 to 43.6 Hz. The single cell RNA transcriptome data of our neuron line supports that the acute effect of Hm1a observed increased the firing activity of GABAergic neurons. We also found that the increased neuron firing rate resulted in the elimination of recurring network bursts present in untreated cultures at similar timepoint. When Hm1a was given to Dravet syndrome model mice by intracerebroventricular injection, both seizure events and epileptiform interictal discharges were significantly reduced (Richards et al., 2018). Further studies are needed to determine if the network synchrony emergent from our neuron preparation represents a model for epileptiform discharges exhibited by Dravet syndrome patients.

HD-MEAs provide a high-throughput approach for reliable recording from human iPSC-derived neuronal cultures at network, single cell, and subcellular resolution (Ronchi et al., 2021). The use of CMOS-based HD-MEAs permitted parallel recording from specific regions of neuronal activity across the array surface, enabling a high-resolution electrical activity map of the neuronal network. The assessment of axonal physiology of the of neurons has historically been difficult to obtain due to technical limitations (Emmenegger et al., 2019). Here we recorded the complete axonal arbors of 1532 neurons. HD-MEAs present the ideal platform for studying axonal physiology due to their high spatiotemporal resolution and ability to record high throughput noninvasively (Bakkum, Frey, et al., 2013; Buccino et al., 2022; Bullmann et al., 2019; Radivojevic et al., 2017). Na_v_1.1 is found at the axon initial segment and nodes of Ranvier of GABAergic interneurons, playing a significant role in axonal physiology (Duflocq et al., 2008; Ogiwara et al., 2007). We confirmed that the Dravet syndrome iPSC-derived neurons had branching axons up to several millimeters in length and showed robust axon potential firing. It is likely due to no myelination, the conduction velocity of the action potential plateaued by DIV 28. CMOS-HD-MEAs are an ideal technology for screening treatments for Dravet syndrome and other neurological disorders where deficits in axonal physiology are implicated. We recorded from the axons of the Dravet patient iPSC-derived neurons over development, demonstrating the feasibility of this technology for high throughput recording of axonal physiology features.

Human iPSC-derived neurons combined with HD-MEA technology provide a high throughput assay platform for furthering the understanding of Dravet syndrome pathophysiology and development of therapeutics. In this study we have recorded at neuronal network, neuron, and subcellular resolution to characterize a patient-derived neuronal culture through development and assay a tool peptide Na_V_1.1 potentiator. Our data supports the continued development of strategies to target Na_V_1.1 rescue to control seizures and ameliorate co-morbidities in Dravet syndrome. The increased inclusion of high throughput electrophysiological readouts of human neurons to drug discovery screening can enable and accelerate the development of novel therapeutic approaches for neurological disorders.

## Methods

### Production of cortical GABAergic neurons from Dravet syndrome patient induced pluripotent stem cells

Human iPSCs derived from peripheral blood mononuclear cells of a Dravet syndrome patient with *SCN1A* Variant 1:1 base pair deletion of T; Nucleotide Position 4522’ Codon Position: 1508; resulting in a frame shift (https://ebisc.org/PFIZi019-A), were differentiated into cortical GABAergic neurons at BrainXell, Inc. following protocol adapted from (Liu et al., 2013).

### Immunostaining

Neurons were thawed and plated on poly-D-lysine (PDL) coated 96-well plates. At DIV 7, media was removed, and cultures washed with Dulbecco’s phosphate buffered saline (D-PBS). Cultures were fixed in 4% paraformaldehyde for 10 min. 2x D-PBS washes were performed and blocking solution added (0.1% Triton X-100, 2.5% donkey serum in D-PBS) for 10 min. Blocking solution was removed, and primary antibodies were added in blocking solution, incubated overnight at 4° C (MAP2 1:1000 Invitrogen #MA5-12826, GABA 1:1000 Sigma Aldrich #A2052). Cultures were washed 2X with D-PBS and blocking performed using 5% donkey serum for 10 min. Blocking solution was removed, and secondary antibodies were added in blocking solution, incubated for 45 min, protected from light (iFluor488 1:1000 AAT Bioquest #16773, iFluor 555 1:1000 AAT Bioquest, #16831, Hoechst 33258 (10 mg/ml, 1:2000), Biotium #40045. Cultures were washed with D-PBS and imaged at 20x. The percentage of Hoescht stained cells expressing MAP2 and GABA was measured from acquired images.

### Characterization of *SCN1A* expression in Dravet patient-derived neurons using qPCR

Dravet syndrome patient iPSC-derived cortical GABAergic neurons were plated with human iPSC-derived cortical astrocytes (BrainXell, Inc., #BX-0600) in a 5:1 ratio on PDL-coated 24 well plate at 300,000 cells per well. At DIV 10, 14, 21 and 28, *n*=3 wells were lysed, and reverse transcription performed using TaqMan Gene Expression Cells-to-C_T_ kit (Life Technologies Corp.) Quantitative PCR (qPCR) was performed using TaqMan assays (ThermoFisher Scientific) for *Scn1a* (Hs0037496_m1) and control gene *TBP* (Hs00427620_m1). The delta-delta CT method was used to calculate fold change of *SCN1A* to DIV 10 expression level (Livak & Schmittgen, 2001).

### Single cell RNA sequencing

A single cell RNA sequencing experiment was performed to understand the cell identity of Dravet patient induced pluripotent stem cell (iPSC) derived neurons. Neurons were thawed following the manufacturer’s protocol and plated in 24-well plate format at 300,000 live cells/well. Base media contained DMEM/F-12 media (0.5x, Gibco), Neurobasal media (0.5x, Gibco), B-27 supplement (1x, Gibco), N-2 supplement (1x, Gibco), GlutaMax (0.5 mM, Gibco), brain-derived neurotrophic factor (BDNF, 10 ng/mL, PeproTech), glial-derived neurotrophic factor (GDNF, 10 ng/mL, PeproTech), and TGF-β1 (1 ng/mL, PeproTech), for a total of 500 mL/well. For seeding media at DIV 0, GABA neuron seeding supplement (1x, BrainXell) was added. At DIV 4, base media plus Day 4 Supplement (1x, BrainXell) and Supplement K (1x, BrainXell) were added at 500 mL/well. Starting at DIV 7, a 50% media change 2x per week was performed using base media. Cells were disassociated at DIV 21 following protocol for mature neuronal culture outlined (Julie Jerber, 2020). Cells in suspension were 89% viable (Countess II) following disassociation. Initial steps of sample QC and cDNA synthesis were conducted at GENEWIZ, LLC./Azenta US, Inc (Waltham, MA, USA). Library preparations, sequencing reactions, and data analysis were conducted at GENEWIZ, LLC./Azenta US, Inc (South Plainfield, NJ, USA).

### Sample QC and Library preparation

Live cell count was accessed using Nexcelom Cellaca Cell Counter along with Nuc blue for all cell nucleus staining and propidium iodide for dead cell nucleus staining. Sample with cell counts > 50,000 and viability > 70% was diluted and loaded onto the Chromium Controller. Loading was performed to target capture of ∼3,000 GEMs per sample for downstream analysis. Sample was processed through the Chromium Controller following the standard manufacturer’s specifications.

Single cell RNA library was generated using the Chromium Single Cell 3’ kit (10X Genomics, CA, USA). The sequencing library was evaluated for quality on the Agilent TapeStation (Agilent Technologies, Palo Alto, CA, USA), and quantified by using Qubit 2.0 Fluorometer (Invitrogen, Carlsbad, CA). Sample was also quantified using qPCR (Applied Biosystems, Carlsbad, CA, USA) prior to loading onto an Illumina sequencing platform. The final library was sequenced at a configuration compatible with the recommended guidelines as outlined by 10X Genomics.

### Data processing and data analysis

Raw sequence data (.bcl files) generated from Illumina was converted into fastq files and de-multiplexed using the 10X Genomics’ cellranger mkfastq command. Subsequent UMI and cell barcode de-convolution was also performed prior to downstream analysis. Mapping to the respective genome and gene expression analysis was performed on Rosalind’s platform. Data was analyzed by ROSALIND® (https://rosalind.bio/), with a HyperScale architecture developed by ROSALIND, Inc. (San Diego, CA). Quality scores were assessed using FastQC (Andrews, 2010). Cell Ranger was used to align reads to the Homo sapiens genome build GRCh38, count UMIs, call cell barcodes, and perform default clustering. In R using the Matrix package, UMAP projection data and feature-barcode matrices outputs from cellranger were converted to csv format which included gene expression data and UMAP projection coordinates for each barcoded cell. Natural log of the expression counts of select genes were projected on UMAP plots to create heatmap plots using R’s ggplot2.

### MEA plating and maintenance of neuronal cultures

Recordings were performed on complementary-metal-oxide-semiconductor (CMOS)-based high density-multielectrode arrays (HD-MEAs) in 6-well plate format (Maxwell Biosystems AG). Two days prior to plating, wells were treated with 1.5 mL of 1% Terg-a-zyme (Sigma Aldrich) solution in deionized water and kept at room temperature for 2 hours to improve hydrophilicity of the array surface. Wells were washed 3x with deionized water and treated with a 70% ethanol solution for 30 minutes for sterilization. Following another 3x wash with deionized water, wells were filled with 2 mL of complete media and kept for 2 days within a 5% CO_2_ incubator at 37°C. On day of cell plating, media was aspirated from each well and 1x wash with deionized water was performed. Arrays were treated with 50 mL 0.1 mg/mL poly-D-lysine (PDL) hydrobromide (Sigma Aldrich) diluted in borate buffer and kept for 3 hours in 5% CO_2_ incubator at 37°C. Wells were washed 3x with deionized water and dried. A secondary plating of Geltrex, 1:100 in DMEM/F12 (Gibco) was applied and kept for 1 hour in 5% CO_2_ incubator at 37°C until aspirated and cells immediately plated. Cortical GABAergic neurons were plated with human iPSC-derived cortical astrocytes from an apparently healthy normal human (BrainXell) at a ratio of 100,000 neurons to 20,000 astrocytes per MEA. Frozen cell vials were thawed following manufacturer’s protocol, combined at 5:1 ratio, and centrifuged in 15 mL conical vial at 160 g to enrich to density of 120,000 live cells / 10 mL. Seeding media: base media plus GABA neuron seeding supplement 1x (BrainXell), astrocyte supplement 1x (BrainXell) and Geltrex 15 mg/mL was prepared fresh on day of cell thaw. 10 mL of cell solutions was placed on center of each MEA and plate placed for 1 hour in 5% CO_2_ incubator at 37°C for cells to adhere. Wells were then filled with 1.5 mL of seeding media, plates sealed with BreathEasy membrane (Sigma Aldrich), and placed in 5% CO_2_ incubator at 37°C for duration of experiment. At DIV 4, media was prepared DMEM/F12 medium (0.25x), neurobasal medium (0.25x), BrainPhys medium (0.5x), B27 Supplement (1x), N2 Supplement (1x), GlutaMax (0.5 mM), BDNF (10 ng/mL), GDNF (10 ng/mL), TGF-β1 (1 ng/mL), Day 4 Supplement (BrainXell), Astrocyte Supplement 1x (BrainXell) and Supplement K 1x (BrainXell) was added at 1.5 mL / well. At DIV 7, a 50% media change was performed with maintenance media: BrainPhys medium (1x), B27 Supplement (1x), N2 Supplement (1x), GlutaMax (0.5 mM), BDNF (10 ng/mL), GDNF (10 ng/mL), TGF-β1 (1 ng/mL). Media changes continued 2x per week for duration of experiment.

### HD-MEA recordings

Each high-density MEA contained 26,400 electrodes organized in 120 x 220 grid within a sensing area of 9.3 x 5.45 mM, and electrode pitch of 17.5 mM (Maxwell Biosystems AG). Configurations of 1024 electrodes could be simultaneously recorded from the array. Recordings were performed weekly from week 1 (DIV 7) till week 6 (DIV 42) at 5% CO2 and 37°C environment (MaxTwo System, Maxwell Biosystems AG). The Activity Scan Assay and Network Assay modules were used from the MaxLab Live software (MaxWell Biosystems AG) with a spike detection threshold of 5.5 standard deviations. For Activity Scan, seven electrode configurations, including 6600 electrodes at a pitch of 35 mm, were recorded for 30 seconds. These recordings mapped the spontaneous activity across the surface of the HD-MEA. For the network recordings, the electrodes with maximum spike amplitude were identified and 3×3 grid of 9 electrodes was recorded around each electrode for a neuronal unit configuration. Each network recording was performed for 5 minutes. Recordings were analyzed using the Batch Analysis Module within MaxLab Live.

### Axon Tracking Assay

The AxonTracking Assay (Maxwell Biosystems AG) was performed to identify single units and measure features of axonal physiology. For each MEA, 50 neurons were identified, and axonal features detected: total axon length, longest axonal branch length, conduction velocity, unit firing rate, spike amplitude at axonal initial segment, and longest latency from axonal initial segment to axon termination.

### Spike sorting

Spike sorting was performed on 5-minute network recordings to identify single neuronal units. The spike sorting pipeline was performed in Python within SpikeInterface (Buccino et al., 2020). Recordings were centered and bandpass filtered from 300-4000 Hz. The SpykingCircus algorithm (Yger et al., 2018) was used for spike sorting with a detection threshold of 6 standard deviations and template width of 5 milliseconds. Sorted units were excluded with a signal-to-noise ratio below 5 and ISI violation rate greater than 0.2.

### Feature extraction

Features were extracted for spike sorted units. These features included single unit waveform features, single unit activity features, and network activity features.

### Single-unit waveform features

The maximum waveform (Figure 4D) was extracted by determination of electrode with maximum amplitude for each unit and averaging recorded unit waveforms.

#### Waveform features

1. **Amplitude**: The voltage amplitude defined as difference between second maximum peak and the minimum (Weir et al., 2014).
2. **Half-width**: The spike half-width was calculated as the width of the spike at half-maximal amplitude of the first maximum (Weir et al., 2014).
3. **Peak latency**: The peak latency was calculated by difference in time between spike minimum occurrence and second maximum peak occurrence (Weir et al., 2014).
4. **Asymmetry ratio**: Asymmetry ratio was calculated by the ratio of amplitude of second maximum peak to amplitude of first maximum peak (Weir et al., 2014).
5. **Trough to peak ratio**: Trough-to-peak ratio was calculated as ratio of amplitude of second maximum peak to amplitude of spike minimum.
6. **Decay**: Decay was calculated by time duration from second maximum peak to crossing zero voltage.
7. **Repolarization slope**: The slope of point of maximum negative EAP to second peak.
8. **Recovery slope:** The slope of second peak to first zero voltage crossing.

#### Unit activity features

9. **Spike frequency (Hz)**: The count of neuronal spiking events divided by duration of recording in seconds.
10. **Interspike interval (ms):** The mean interspike interval for each neuronal unit for duration of recording.
11. **Intraburst frequency (Hz):** Neuronal bursting events were detected using the logISI method (Pasquale et al., 2010). The intraburst frequency of neuronal firing was determined by averaging the frequency of firing within the bursts of each individual neuron during the recording.
12. **Burst frequency (bursts/min):** Frequency of neuron bursting was determined by count of bursts within recording divided by duration in minutes.
13. **Burst duration (sec):** For each neuron, the duration of each detected unit burst.
14. **Number of spikes per burst:** For each neuron, the number of spikes within each detected unit burst.
15. **Intraburst interspike interval (ms):** For each neuron, the mean interspike interval for each detected unit burst.

#### Network activity features

16. **Burst frequency (bursts/min)**: Network bursts from spike sorted units were detected using the ISI_N_ method, with *N*=10. (Bakkum, Radivojevic, et al., 2013). Frequency of network bursting was determined by count of bursts within recording divided by duration in minutes.
17. **Burst duration (sec):** The mean duration of network bursting events for a recording.
18. **Spikes per burst:** The number of neuronal spikes within a network burst.

### Pharmacology experiments

At DIV 42, HD-MEAs were treated with pharmacological agents to assess neuronal responses. Following a 30 second period of baseline recording in the neuronal network configuration, each drug diluted in vehicle (PBS), was added to the HD-MEA media in a 10 μL volume to achieve final drug desired concentration. A recording then continued for 5 minutes to assess neuronal firing responses. The GABA_A_R receptor agonist muscimol (10 μM, Tocris, 0289) was used to assess for functional GABA_A_R signaling. The Na_V_1.1 potentiator δ-theraphotoxin-Hm1a (Hm1a, Alomone Labs, #STH-601), used at 100 and 500 nM. A total of *n*=3 HD-MEAs was recorded from for each drug and vehicle (PBS). Recordings were spike sorted like above and split into 30 seconds of baseline activity and 30 seconds of post-drug activity initiated at treatment start time. Instantaneous spiking frequency was determined by averaging spikes per neuron in 2 second bins. To assess the effects of the treatments on the individual neuron firing rate, the baseline and drug periods were split into 5 second bins of averaged firing rate. The maximum firing rate was compared for each neuron between baseline and treatment to determine a Δ frequency in hertz. Neuron burst firing was detected using the LogISI algorithm (Cotterill et al., 2016; Pasquale et al., 2010).

### Confocal imaging of immunostained HD-MEAs

Following recording experiments at DIV 42, media was removed from HD-MEAs and following a PBS wash, wells were fixed for 10 minutes with 4% paraformaldehyde solution. Cells were permeabilized with a 0.1% Triton-X-100 (Sigma Aldrich) solution for 15 min at room temperature. A blocking solution was then added with 0.1% Triton-X-100 and 3% bovine serum albumin (Invitrogen). An antibody solution of MAP2 (1:2500, Invitrogen #PA1-10005), GAD-67 (1:200, Invitrogen #MA5-24909), and V-Glut-1 (1:1000, Invitrogen #48-2400) was added and incubated on shaker for 1 hour at room temperature. Wells were washed 3x with PBS, and a secondary antibody solution added containing AF-647, AF-555, and AF-488 (1:1000, Invitrogen #A32933, #A32732, #A32766). Wells were incubated for 1 hour in the dark. A PBS wash was performed and Hoescht staining solution added (1:2000 in PBS, ThermoFisher) and wells incubated for 5 minutes on shaker. Following a 3x PBS wash, images were obtained on a Zeiss LSM880 upright confocal point scanning microscopes with a 20x NA 1.0 water dipping lens. To accommodate the large MEA sensor area and large cell clusters, the tiling functionality with Z-stacks were used to capture this volume in its entirety (∼150um in Z, 2.5×5 mm in X/Y). This volumetric image was then stitched and collapsed into a single 2D plane.

### Code

All code for spike sorting and electrophysiological analysis was performed using Python 3. Analysis of scRNA sequencing data was completed using R.

### Statistics

Values are mean values ± standard error of the mean ± SEM unless indicated. All statistical tests were performed with Prism 9 (Graphpad, La Jolla, CA, USA). Multiple groups were compared using the Kruskal-Wallis test, followed by the Dunn-Sidák multiple-comparison test. Comparisons between two values were performed using the Mann-Whitney test. Significance was considered at *p* values <0.05.

### Data Availability

The scRNA-seq sequencing data and processed files were deposited in the GEO database under accession code GSE245113.

## Acknowledgements

The Pfizer Cellular and High-Content Technology Center for confocal imaging of HD-MEAs.

